# Why would parthenogenetic females systematically produce males who never transmit their genes to females?

**DOI:** 10.1101/449710

**Authors:** Manon Grosmaire, Caroline Launay, Marion Siegwald, Marie-Anne Félix, Pierre-Henri Gouyon, Marie Delattre

## Abstract

**Summary:** In pseudogamous species, females use the sperm of males from another species to activate their oocytes and produce females, without using the sperm DNA. Here we report a novel reproductive strategy found in the pseudogamous nematode *Mesorhabditis belari*, which produces its own males at low frequency. We find that the 8% of *M. belari* males are necessary to fertilize all oocytes but pass on their genes only to males, and never to females. Thus, the production of males has no impact on the genetic diversity of females. Using game theory, we show that the production of males at low frequency constitutes an efficient strategy only if sons are more likely to mate with their sisters. We validate this prediction experimentally by revealing a mating preference between siblings. We uncover the remarkable reproductive strategy of parthenogenetic females that pay the cost of producing males while males do not spread their genes.

## Results

Sperm-dependent parthenogenesis, also called pseudogamy, is a reproductive strategy in which females use the sperm of males, usually from another species, to activate their oocytes. Later, the sperm DNA does not participate to the development of the zygote, leading to 100% females in the progeny (1, 2). In the nematode family Rhabditidae, a very special type of pseudogamy is found within the *Mesorhabditis* genus, where the sperm required to activate the oocytes does not come from males of other species: in pseudogamous species *of Mesorhabditis*, both females and males are produced. This unique reproductive strategy is characterized by a low frequency of males in the population (below 10%). Following up on initial descriptions by Belar in the 1920’s (3) and Nigon in the 1940’s (4), we recently reisolated strains of pseudogamous *Mesorhabditis belari* (initially named *Rhabditis belari (4, 5)*) and revisited the results obtained by Belar and Nigon. Here we show that females arise strictly by gynogenesis, without incorporating the male DNA into their genome, while males arise strictly by amphimixis of the parental genomes. Thus, the genes of males are never passed on to females in this species. From the point of view of the gene pool of females, the fitness return on males is null. We next ask why females that have the ability to reproduce by gynogenesis would still produce males which do not spread their genes to females. Using game theory, we find that the production of less than 10% of males constitutes an evolutionary stable strategy to optimize the number of oocytes that can be fertilized, as long as the males have a mating preference for their sisters. Transient or cyclical production of males by parthenogenetic females have been described in several species (6). In all cases, the production of a sexual form leads to exchange of genetic material, thus introducing genetic diversity in an otherwise clonal population (7–11). To our knowledge, the reproduction of pseudogamous Mesorhabditis nematodes is a novel and unexpected strategy in which parthenogenetic females produce a low percent of males while males do not provide any genetic diversity to the females.

### *Mesorhabditis belari* females produce gynogenetic and amphimictic embryos

We chose to focus on *M. belari* JU2817 (Orsay, France) as our reference strain (see Methods). In our laboratory conditions, females of this strain produced approximately 8% of males during their entire life (mean=7,72%, standard deviation=2,2, count of 3743 F1s, see Methods) (Table S1). We also confirmed that females could not reproduce in the absence of males. Last, we found a level of embryonic lethality of only 5% (see Methods), ruling out the possibility that the low percentage of males was due to high lethality of male embryos.

In *M. belari* females, as in other Rhabditids nematodes, oocytes are arrested in prophase of meiosis I and meiosis resumes after fertilization (12). Using DIC recordings and staining of DNA and microtubules from fixed samples, we found, as described by Belar and Nigon (3, 4), 2n=20 chromosomes in female oocytes (Figure S1) (4). We also found that females produced two types of embryo (Figure 1A and 1B). (i) The most common class were gynogenetic embryos (n= 227 out of 258 fixed embryos), which extruded a single polar body and displayed a single pronucleus, containing 20 chromosomes before the first embryo mitosis (Figure S1); in this case the oocyte followed a single round of meiotic division and the male DNA entered the oocyte cytoplasm but did not decondense; as suggested by Belar, we found that despite the absence of sperm DNA decondensation, the centrosomes that were provided by the sperm formed robust asters that eventually contacted the female pronucleus to establish the first mitotic spindle (ii) A few % of the embryos (n=31 out of 258 fixed embryos) were amphimictic: two polar bodies were observed, two pronuclei containing 10 chromosomes each were formed before mitosis (Figure S1) and asters were seen associated with the male DNA; in this case, the oocyte followed two rounds of meiotic divisions and the haploid male DNA decondensed and mixed with the haploid female DNA. Both categories of embryos were diploid, because the lack of paternal DNA was compensated by the lack of one female meiotic division in gynogenetic embryos.

**Figure 1.**
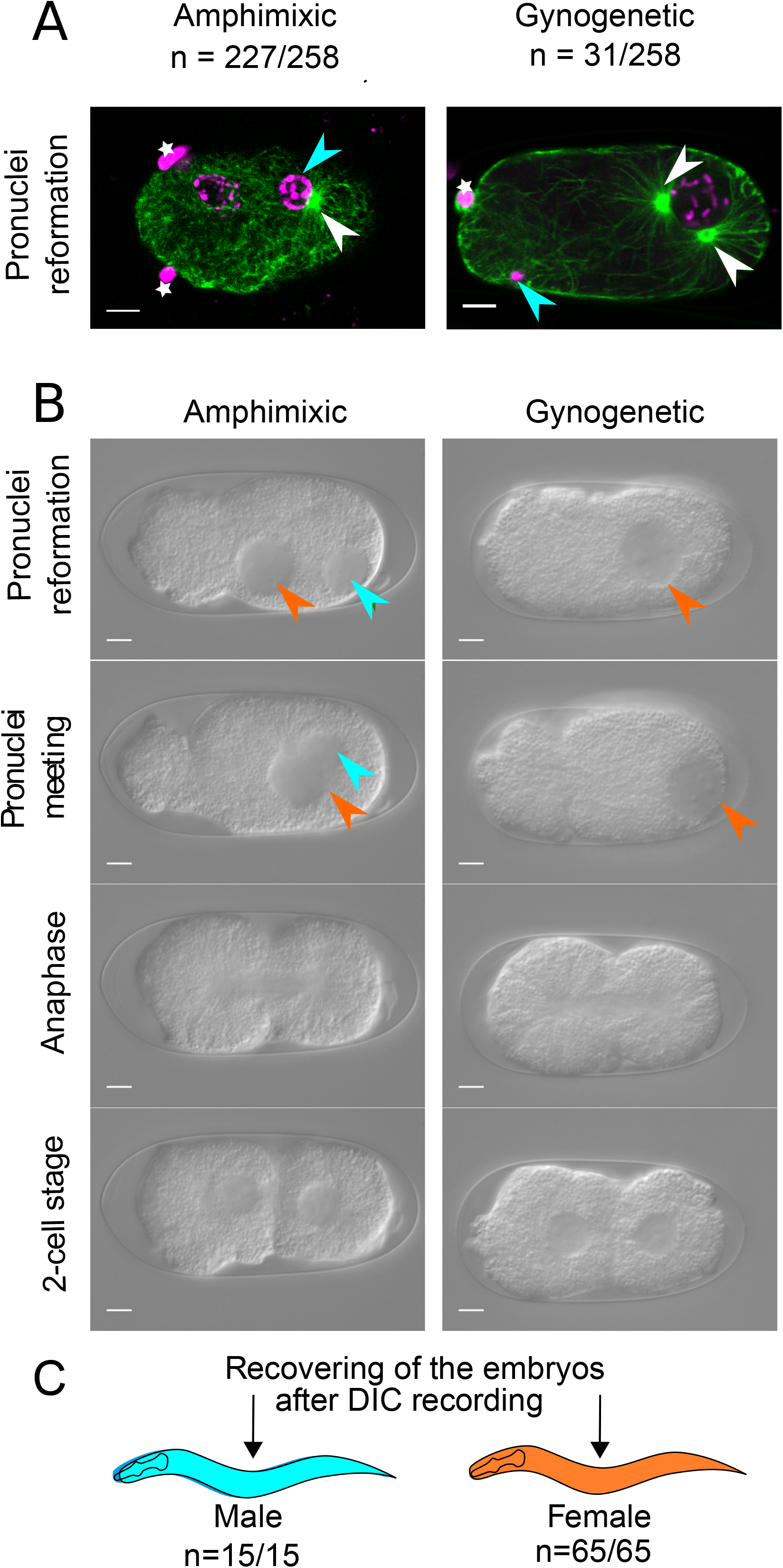
Two types of embryo are produced by *Mesorhabditis belari* females. (A) Images from a representative amphimictic and a gynogenetic embryo during the reformation of the pronuclei fixed and stained with an anti-tubulin antibody (in green) and with Hoechst to label DNA (in magenta). White arrowsheads point towards the centrosomes. Note that in the amphimictic embryo, only one aster is visible on this focal plane. Polar bodies are shown with a white star and male DNA is shown with a blue arrowhead. Out of 258 one-cell embryos scored during pronuclei reformation, only 31 showed two pronuclei. In gynogenetic embryos, the male DNA stays highly condensed, the female pronuclei has captured the centrosomes and only one polar body is visible. Scale bar is 5 µm. (B) Still images from DIC recordings of representative amphimictic or gynogenetic embryos of *M. belari*. The male (blue arrowhead) and female (orange arrowhead) pronuclei are followed from fertilization until the first cell division. Scale bar is 5 µm. (C) *M. belari* embryos were sorted based on the number of polar bodies and pronuclei observed on DIC recordings, as shown in (B). 65 gynogenetic embryos always gave rise to 65 females (in orange), while 15 amphimictic embryos gave only males (in blue).

Regardless of the status of the male DNA, the sperm is required to activate all the oocytes and to provide centrosomes to the zygote in the *M. belari* species.

### The DNA of *M. belari* males is only transmitted to sons

After observing this first cell division by DIC time-lapse imaging (Figure 1B), we recovered embryos of each category and let them develop on an agar plate. We found that 65 out of 65 gynogenetic embryos (harboring a single pronucleus) developed as females. In contrast, we found that 15 out of 15 amphimictic embryos (showing two pronuclei) developed as males (Figure 1C). Instead, Nigon had found that 7 out of 10 of the amphimictic embryos gave rise to males and 3 to females. Based on this result, he suggested that amphimictic embryos produced 50% of males and 50% of females, as in regular amphimictic species (4).

To confirm our finding that amphimictic embryos did not produce females, we took advantage of a single-nucleotide polymorphism (SNP) within the RNAP2 locus of different strains of *M. belari* to genotype males and females. We crossed females from the JU2817 strain (with a G/G genotype) with males from the JU2859 strain (with a C/C genotype) and vice versa. We genotyped the F1s from different crosses and found that, irrespective of the cross direction, all F1 females had the same allele as their mother (n=539), while the F1 males were all heterozygotes C/G (n=49) (Table 1). This result indicates that *M. belari* female individuals arise strictly by gynogenesis, while amphimixis produces only males. Because one-cell embryos with two pronuclei are difficult to distinguish from 2-cell stage embryos under low magnification, we believe that Nigon missorted three female embryos. However, we cannot exclude that the strain that was used by Nigon behaved differently as those that we had recently isolated.

**Table 1.**
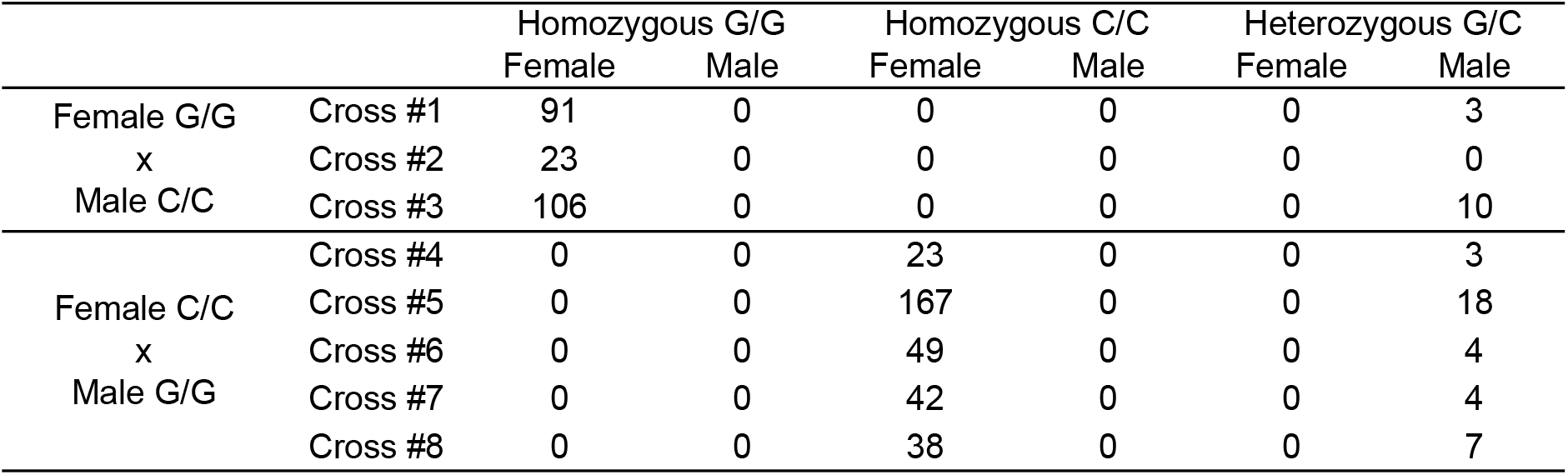
Females come from gynogenesis and males from amphimixis. *M. belari* JU2817 are homozygous G/G at a RNAP2 locus, while *M. belari* JU2859 are homozygous C/C at the same locus. JU2817 females were crossed to JU2859 males and vice versa (crosses 1-3 or crosses 4-8, respectively) and their entire progeny was genotyped. For all crosses, while males always presented a heterozygous combination of the parental alleles, females always harboured the genotype of their mothers.

Our results imply that *M. belari* JU2817 females produce males whose DNA is never transmitted to females. Our results also suggest that the male genome is diluted in their sons at each generation with the mother genome, thus in principle converges with the female genome. However, males could transmit preferentially the alleles of their father to their sons, thus preventing this dilution. To test this hypothesis, we genotyped the sons of heterozygous males. To this end, we first generated heterozygote males C/G at the RNAP2 locus by crossing JU2817 G/G females to JU2859 C/C males. Heterozygote males were then crossed to JU2859 C/C females. We found that out of 48 F1 males, 24 were C/C and 24 were C/G. Thus, heterozygote males transmitted the RNAP2 alleles of their parents at the same frequency. This result suggests that there is no global bias in the transmission of the paternal genome in *M belari*, although we cannot exclude that some loci and/or chromosomes are subjected to a male meiotic drive. Overall, we propose that male genes are progressively converging with the female genes in males, and because they will never be found in females, there is no selection on most of the male genes. From the point of view of the gene pool of *M. belari* females, the fitness return on males is 0.

### A male proportion of 8% is sufficient to maximize the brood size of females

We next asked whether a proportion of 8% males in the population was sufficient to obtain the largest possible brood size, either the number of possible male mating or the number of transferred sperm cells being potentially a limiting factor. In other words, we asked whether this low percentage of males was optimal for the fitness of the population. To answer this question, we crossed 20 females with an increasing number of males and counted the total number of progeny that were produced during the reproductive period (Figure 2 and Table S2). We found that batches of 20 females that had been crossed to a single male produced in average approximately 490 descendants, while batches of 20 females that had been crossed to two males reached an average of 800 descendants. Importantly, further increasing the number of males per females did not increase the mean brood size of females. This result demonstrated firstly that the mean production of a single female could not exceed 40 embryos, even when males and sperm cells were in excess. In agreement with this finding, we showed that the mean number of sperm cells stored in the spermatheca of females ranged from 64 to 226 sperm cells (mean = 134.72 +/- SD = 47.52, n=25 females), largely exceeding the number of embryos produced during the reproductive period of females. Secondly, our result showed that a single male was not sufficient to fertilize and transfer enough sperm cells to 20 females and therefore, male percentage was a limiting factor in the population. However, two males for 20 females, i.e. 9% of males, were already sufficient to obtain the maximum brood size. This result strongly suggested that a single male could produce between 400 and 800 sperm cells during its reproductive period. Overall, we concluded that the production of 8% amphimictic (male) embryos was sufficient to maximize the brood size of *M. belari* JU2817 females.

**Figure 2.**
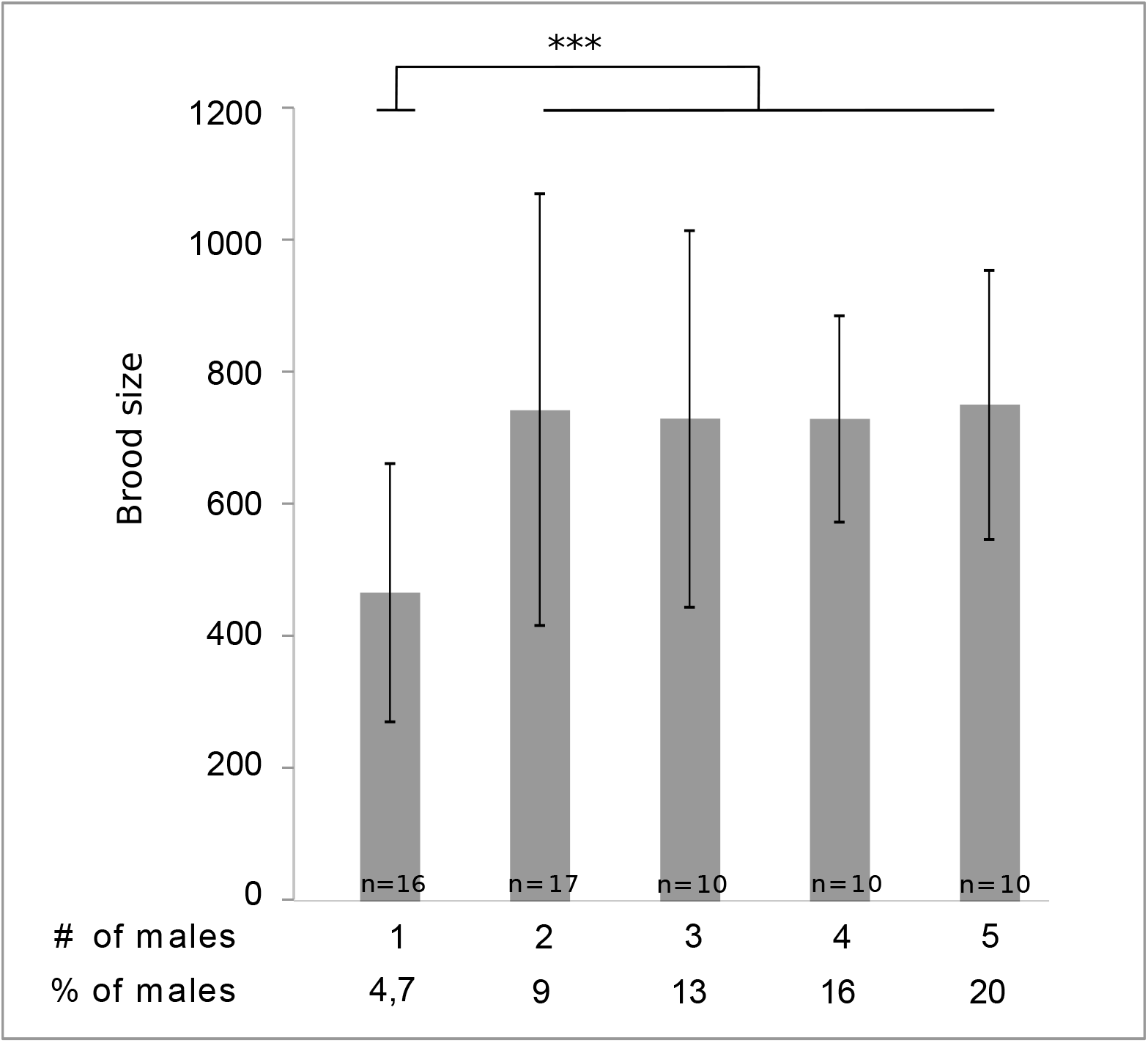
The percentage of males in the population is sufficient to maximize the brood size of females. 20 females were crossed with a different number of males (from 1 to 5, corresponding to 4.7 to 20 % of males) and left on plates until the end of the female reproductive period. The total number of embryos produced (Y axis) was counted for *n* crosses per condition. The mean value and standard error are shown for each condition. Corresponding numerical values are shown in Table S2.

### Evolutionary stable strategy for the sex ratio in *M. belari*

The interesting peculiarity of this system is that the genes present in a male will never be in a female again. From the point of view of the genes of females, the fitness return from sons is equal to 0. Thus, production of sons might be of interest for a female only if its sons mate with its daughters (to ensure that a maximum of eggs is fertilized) and do not mate unrelated lineage of females (to which they do not transmit any DNA). Because gynogenetic females could theoretically be fertilized by males of another species, we asked under which conditions should *M. belari* gynogenetic females produce their own males and in which proportion? To answer this question, we developed a game theory model. We show below that the *M. belari* reproductive strategy cannot be maintained in large populations under panmixis, suggesting that *M. belari* females should be isolated from the others either by physical distance (i.e the descendants do not move much before mating and mate with their siblings) or by their behavior (i.e males prefer mating with their sisters rather than with unrelated females).

In the model, the proportion of sons produced by a female will be called the sex ratio and noted *x*. The basic hypothesis of the model is that the progeny of a female constitutes a patch (or deme) of size *k*, which exchanges a fraction of *m* migrants with other demes (here this migration rate can also be interpreted like a rate of mating with unrelated females if the isolation is behavioural rather than geographical). Males and females are supposed to migrate equally. The fitness *W*(*x, x*^*^) of a female playing a sex ratio *x* in a population where the others play a sex ratio *x*^*^ is computed as the number of fertilized eggs that her daughters produce. The evolutionary stable strategy (ESS) will be found by searching the value of *x*^*^ such that *∀ x* ≠ *x*^*^, *W(x, x) < W(x, x^*^*). The function *W(x, x^*^*) reaches its maximum when *x* = *x*^*^.

We first developed an analytical model (i). We explored the ESS sex ratio considering that the males are produced in order to ensure that at least one male will be present in the deme of the daughters, supposing that one male will be enough to make all eggs fertile. Second, we explored the ESS sex ratio using computer simulation (ii), taking into account that each male produces a finite number of spermatozoa.

#### (i) The analytical model: unlimited sperm

Within the deme constituted by the progeny of female playing *x*, the sex ratio after migration is: 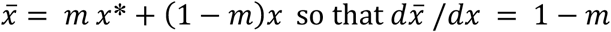

The probability that there is no male in the deme is 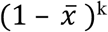 and in the other demes (their number is supposed to be much larger than 1 so that the influence of the sex ratio of female *x* can be neglected once her progeny is dispersed) is (1 - *x*^*^)^k^.

The fitness of female *x* is then:

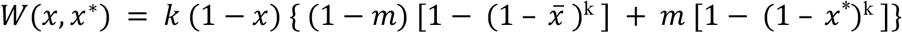

Which leads to an evolutionary stable proportion of females:

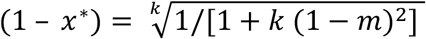

In a panmictic population with infinite migration (*m*=1), the stable sex ratio is equal to 0, because a female has no advantage to produce sons if they mate mostly with the daughters of unrelated females without passing on their DNA. This would secondarily lead to population extinction. However, with decreased migration between demes, the stable sex ratio of the population increases and females do stably produce sons. We found that in order to explain the observed sex ratio of 0.08, it is necessary to suppose a migration rate of less than 0.4 between demes. The curve being relatively flat, migration rates between 0 (no migration) and 0.4 are compatible with the observed sex ratio (Figure 3A).

**Figure 3.**
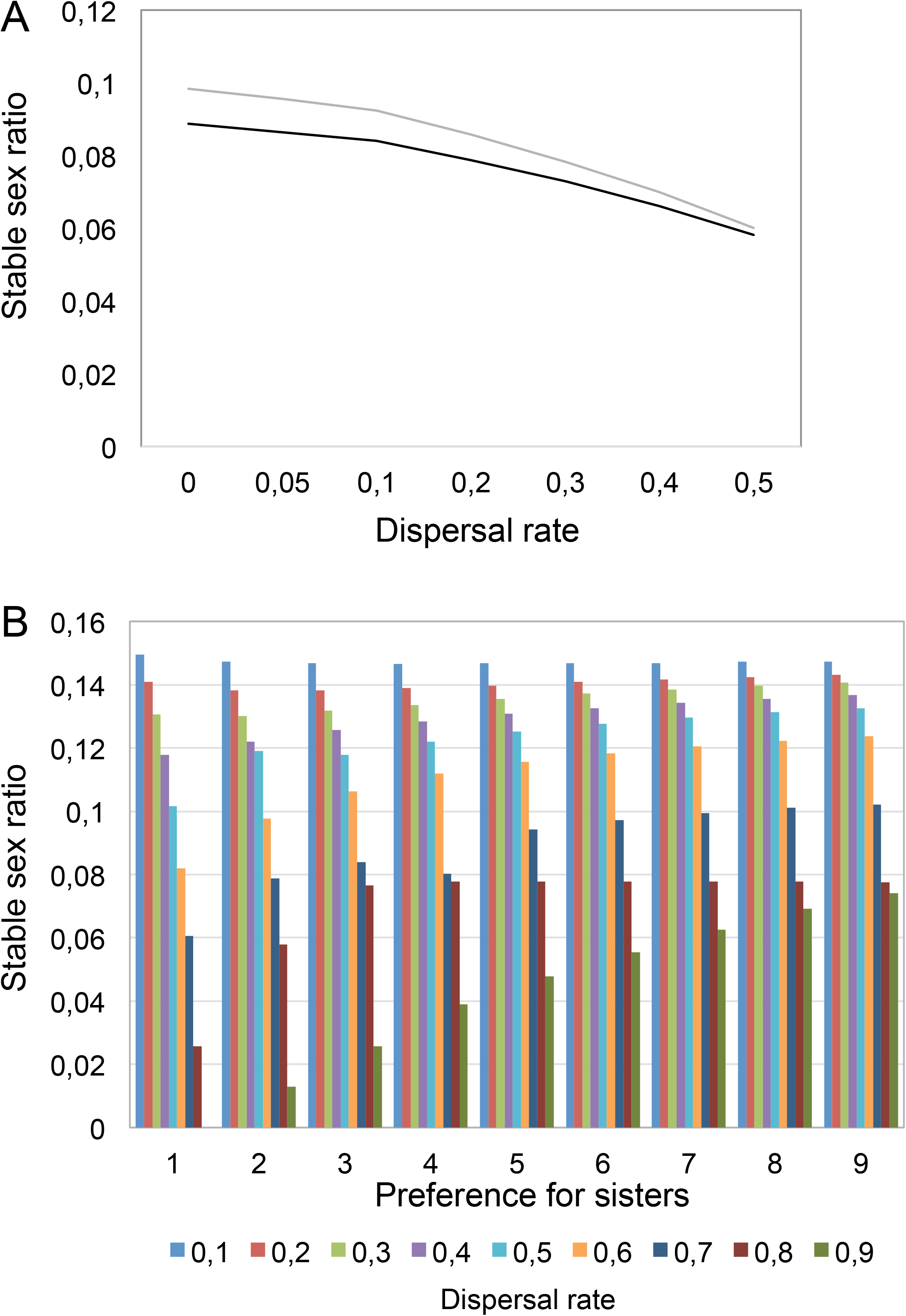
Evolutionary stable sex ratio calculations indicate that a non-zero male proportion arises from a low dispersal rate and/or a high preference for sib-sib mating. (A) The evolutionary stable sex-ratio (proportion of males) is shown as a function of the dispersal rate between demes with 40 individuals per deme and 400 spermatozoa per male, when the sperm is limiting (gey curve) or not limiting (black curve). In a panmictic situation (dispersal rate of 1), no males are produced. Decreasing the dispersal of individuals between demes allows the production of a low percent of males. (B) Influence of the preference for sib-sib mating in the model with sperm limitation, with 40 ovules produced per female and 400 spermatozoa per male. For a strong dispersal rate, the high mating preference between siblings allows the maintenance of few males in the population.

#### (ii) The computer simulation model: limited sperm

In order to take into account that a single male cannot fertilize an infinite number of oocytes (Figure 2A), we simulated the different probabilities of having a certain number of males per deme. Each male produces a limited number of sperm cells, noted *n*. Using simulations, we found the same dependency of sex ratio on migration, although at a given dispersal rate, limiting sperm increases slightly the ESS proportion of males (Figure 3A). For example, a sex ratio of 0.08 is explained by a dispersal rate of 0.15 if the sperm is not limiting, and of 0.3 if sperm limitation is taken into account.

We next distinguished between dispersal and mating preference of males for their sisters. Preference of males for their sisters (ranging from 1= no preference, to 9= prefer their sisters 9 times more often) had a strong impact on the stable sex ratio (Figure 3B). For instance, even if the dispersal rate was very high (0.9), the sex ratio could reach 0.08 if males chose their sisters as a mating partner 9 times more often than unrelated females, while in the absence of mating preference the sex ratio dropped to 0. Hence, *M. belari* females have an advantage of producing males only if the males are more likely to mate with their sisters, either because of a lack of dispersal, or by an active choice of males or females towards their siblings.

### *M. belari* favors sib-sib mating

We next experimentally tested the mating preference of *M. belari* individuals. We had previously observed that in our laboratory conditions, crosses between few males and females that had been isolated randomly from a mixed population were not yielding any progeny in about half of the crosses. We thus tested the possibility that males or females would preferentially mate with individuals that were born from the same mother, and may not yield any progeny when their partners were born from another female. To this end, we isolated single gravid females from the *M. belari* JU2817 strain on plates and performed crosses between 5 females coming from one mother with 2 males coming from the same mother (siblings) or with 2 males coming from another mother (non-siblings). We performed 50 crosses of each category and repeated the experiment 3 times. We found that 117 out of 150 sib-sib crosses were successful, which was statistically higher than the 73 out of 150 successful crosses obtained from non-siblings (p<0.05, χ^2^ test) (Figure 4 and Table S3). These results demonstrated that, as suggested by the mathematical model, *M. belari* individuals prefer mating with their siblings.

**Figure 4.**
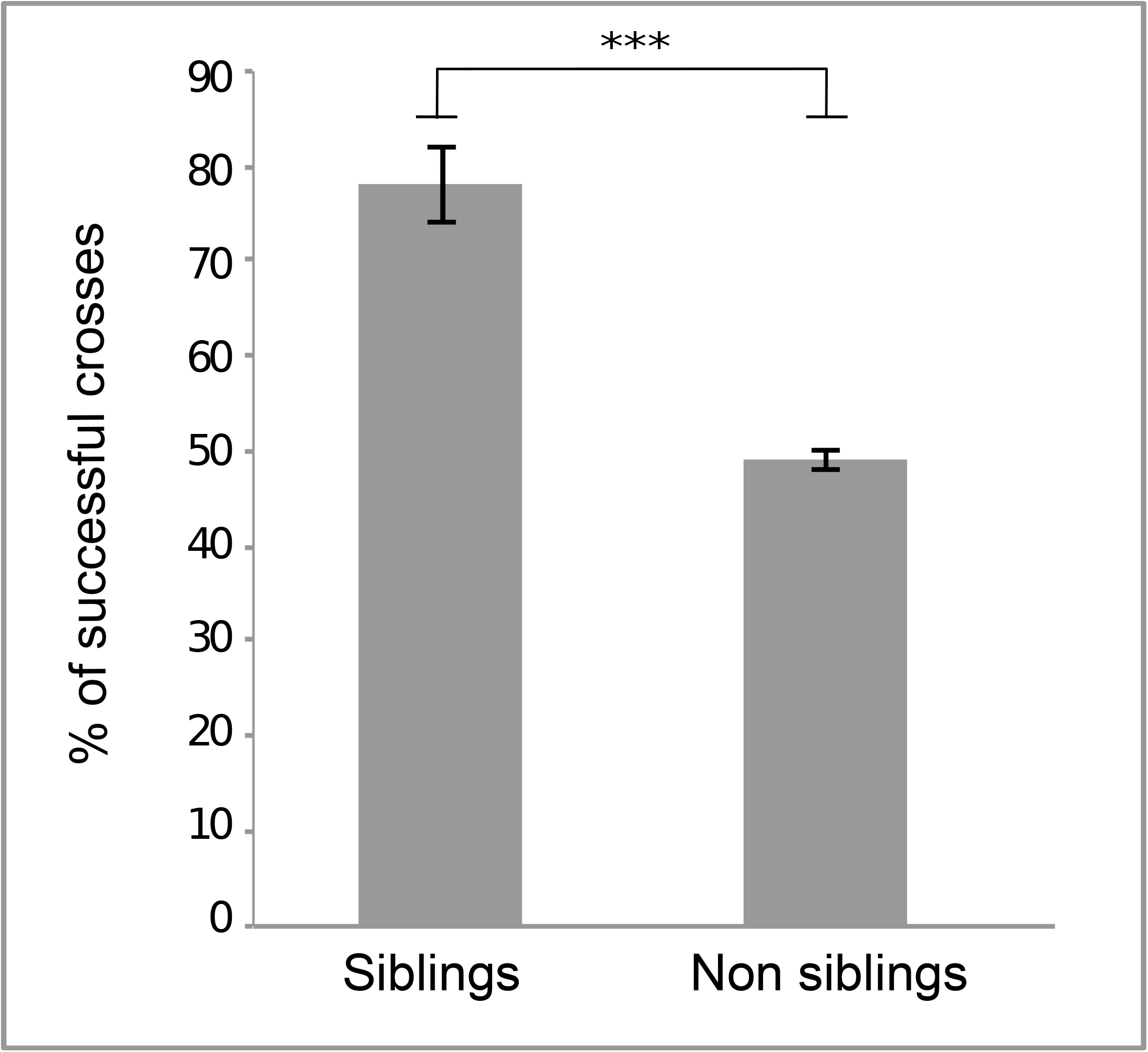
Sib-sib mating preference. We performed 50 crosses between siblings and 50 crosses between non-siblings and counted the % of crosses that gave a progeny (Y axis). The experiment was repeated 3 times. The mean value and standard error are shown for each condition. Corresponding numerical values are shown in Table S3.

## Conclusion

While pseudogamous species can cope with the lack of paternal DNA, they still rely on a sperm cell to activate their oocytes and most likely to inherit centrosomes for embryo development. Indeed, except for some strict parthenogenetic species in which centrosomes form *de novo* after egg activation, the centrosomes are provided by the sperm at fertilization in all animal species that have been studied so far (13). Due to these developmental constraints, pseudogamous females must spend energy for the search of males from another species (1, 2, 14). We have shown that upon identical developmental constraints, the reproductive strategy of *M. belari* consists in producing just enough of its own males. Such strategy may avoid competition for males with females of another species. However, because these males do not transmit their DNA to females, there is no direct advantage to a female (or to her genes) in producing males if they do not enhance their genetic diversity through crosses and if they fertilize mostly the daughters of others without passing on their genes. The only advantage for a sperm-dependent parthenogenetic female to regularly produce males is to ensure that a maximum of its oocytes will be fertilized. We found that because there is a strong mating preference between siblings, the production of less than 10% of males by parthenogenetic females constitutes an efficient reproductive strategy.

Among the known pseudogamous species that use the sperm from another species to reproduce, the production of few males by the pseudogamous females themselves has not been systematically ruled out. In some cases, the existence of rare males has even been reported but interpreted as abnormal events (15, 16). Further investigations will therefore be needed to find out whether convergent evolution as led to the emergence of the remarkable reproductive strategy found in *M. belari* outside of the Mesorhabditis genus.

To our knowledge, the production of males by parthenogenetic species has been systematically interpreted as developmental errors (because they males are found very rarely) (17) or has been associated with the advantage of mixing parental genomes (6). Thus, cyclical parthenogens combine the advantages of both sexual (mixing genomes) and asexual (clonal expansion) reproduction (7–11). The case of *M. belari* shows however that the production of sexual forms cannot be systematically interpreted as a strategy to enhance the genetic diversity of the species.

## Material and Methods

### Strain isolation and maintenance

Nematodes were isolated from soil and rotting vegetal matter and were identified as *Mesorhabditis belari* based on their morphology, geographic occurrence, crosses between strains and sequencing of the ITS2 locus (Genbank accession number MH142258). Strain JU2817 was started from one gravid female isolated from a rotten apple on 20 Oct 2014 in Orsay, France (GPS 48.70200, 2.17282). Strain JU2859 was started from one gravid female isolated in a soil and leaf litter sample collected on 6 Jun 2015 in the Botanic Garden, Cambridge (UK) (GPS 52.19255, 0.1281). We chose JU2817 as our reference strain.

Strains were maintained on Nematode Growth Media and *E. coli* OP50 at 20°C, following the procedure adopted for the maintenance of *C. elegans (18)*. Their life cycle takes around 7 days at 20°C. Males are morphologically distinguishable from females because they are much smaller, have a reduced motility and a characteristic male tail.

### Measurement of embryonic lethality

A single gravid female was isolated and allowed to lay eggs for 24 hours. We counted the number of laid eggs, and 4 days later the number of larvae. We scored 1005 embryos from 79 females and found 50 dead embryos, corresponding to approximately 5 % embryonic lethality.

### Sex ratio measurements

As previously described for *C. elegans*, we found that *M. belari* larvae can be synchronized after an L1 arrest. Briefly, a mixed population of worms was collected in M9 and axenized by bleaching. Embryos were protected by their eggshell and resisted to the treatment. After hatching on plates without food, all worms were arrested at the L1 stage. After two days, L1s were fed with bacteria and restarted their development synchronously. Before they reached adulthood, virgins were isolated. Crosses between 5 virgin females and 2 virgin males were performed on individual plates. Because *M. belari* females lay eggs during approximatively 10 days, parents were transferred to a fresh plate every two days, to avoid the mixing of F1 and F2 generations. We noticed however that frequent transfer of females reduced the size of the brood compared to females that were maintained on their original plate. After 4 days of development, when sexes were distinguishable, the number of F1 males and females was counted. We performed 38 crosses and obtained 3743 F1s, including 289 males, corresponding to a proportion of 7,72 % males, with a standard error of 2,2 (Table S1).

### Measurement of brood size

From a synchronized population of worms (see above), 20 virgin females were isolated and 1 to 5 males were added onto each plate. In order to evaluate the entire brood per female, laid embryos were counted and eliminated every day until the end of the reproductive period (Table S2).

### Preference for sib-sib mating

From a plate of worms from the JU2817 strain, gravid females were isolated on single plates and let to lay eggs. When F1 virgin females and males were distinguishable, they were isolated and crossed with males or females from the same mother (siblings) or from another mother (non-siblings). Five females were crossed to two males and the number of crosses that gave rise to embryos was counted (Table S3).

### Embryo recording

Gravid females were placed into a watch glass containing M9 and cut with a razor blade to release embryos. Embryos were next mounted in M9 on a 2% agarose pad between slide and coverslip, observed on a Carl Zeiss Axioimager A1 microscope with a 100X DIC lens (NA=1.4) and recorded using the time-lapse module of a digital Kappa Camera (DX4-285FW). After recording of the first division, embryos were sorted based on the number of pronuclei and recovered on agar plates. The embryos were allowed to hatch and develop until males and females could be distinguished.

### Embryo and gonad stainings

To stain embryos and spermathecae, gravid females were cut on a polylysin-coated slide in a drop of 0.5 × M9 (diluted 1:1 in H2O). We fixed embryos using a freeze-cracking method, as previously described for *C. elegans* and other nematode species (19). Embryos were fixed during 15 min in −20°C methanol. A mouse anti-tubulin antibody (Sigma-Aldrich DM1a 1/200) and a donkey anti-mouse secondary antibody (DyLight 488, Jackson Immunoresearch 1/1000) were incubated at room temperature during 45 min. Slides were finally incubated during 5 min in 1μg/ml Hoechst 33342 (Sigma-Aldrich) to reveal DNA. Labeled embryos were observed with a confocal microscope (LSM710, Zeiss). Z-stacks of embryos were acquired every 0,29 µm. To stain the spermatheca, females were cut to release the gonad and a drop of paraformaldehyde 3% was added onto the samples for 5 minutes. Next, a coverslip was placed on the sample and frozen on a cold metal block. After removal of the coverslip, slides were immersed into methanol at −20°C and then rinsed and stained with Hoechst, following the same procedure as for embryo staining. Sperm cells inside the unique spermatheca of *M. belari* females were counted manually using an epifluorescent AxioImager A2 Zeiss microscope.

### Worm genotyping

Sanger sequencing revealed that two *M. belari* strains (JU2817 and JU2859) had an identical sequence of the RNAP2 gene except for a single single-nucleotide polymorphism (SNP). At this locus, the JU2817 strain displays a homozygote G/G genotype, while JU2859 has a C/C genotype. From synchronized worm populations (see above), virgin females and males from each strain were mated on single plates. PCR on whole worms were performed to genotype the F1s. We performed pyrosequencing of the F1s from JU2817 females crossed to JU2859 males. We used High Resolution Melt (HRM) sequencing to genotype the F1s from a cross between JU2817 males and JU2859 females. DNA was prepared using a Qiagen Type-it HRM PCR kit and a Qiagen Rotorgene following the instructions of the supplier.

### Models

The analytical model is shown in Appendix 1. The simulation was written in Python and is available upon request.

## End notes

This work has been supported by a grant from the PEPS CNRS (from 2014 to 2015) and by a grant fellowship to MG from the French ministry of research (from 2014 to 2018). MD and MAF designed the experiments. PHG designed the model and performed the simulations together with MS. MG, CL and MD performed the experiments. MG, MAF, PHG and MD wrote the manuscript.

**Figure S1.**
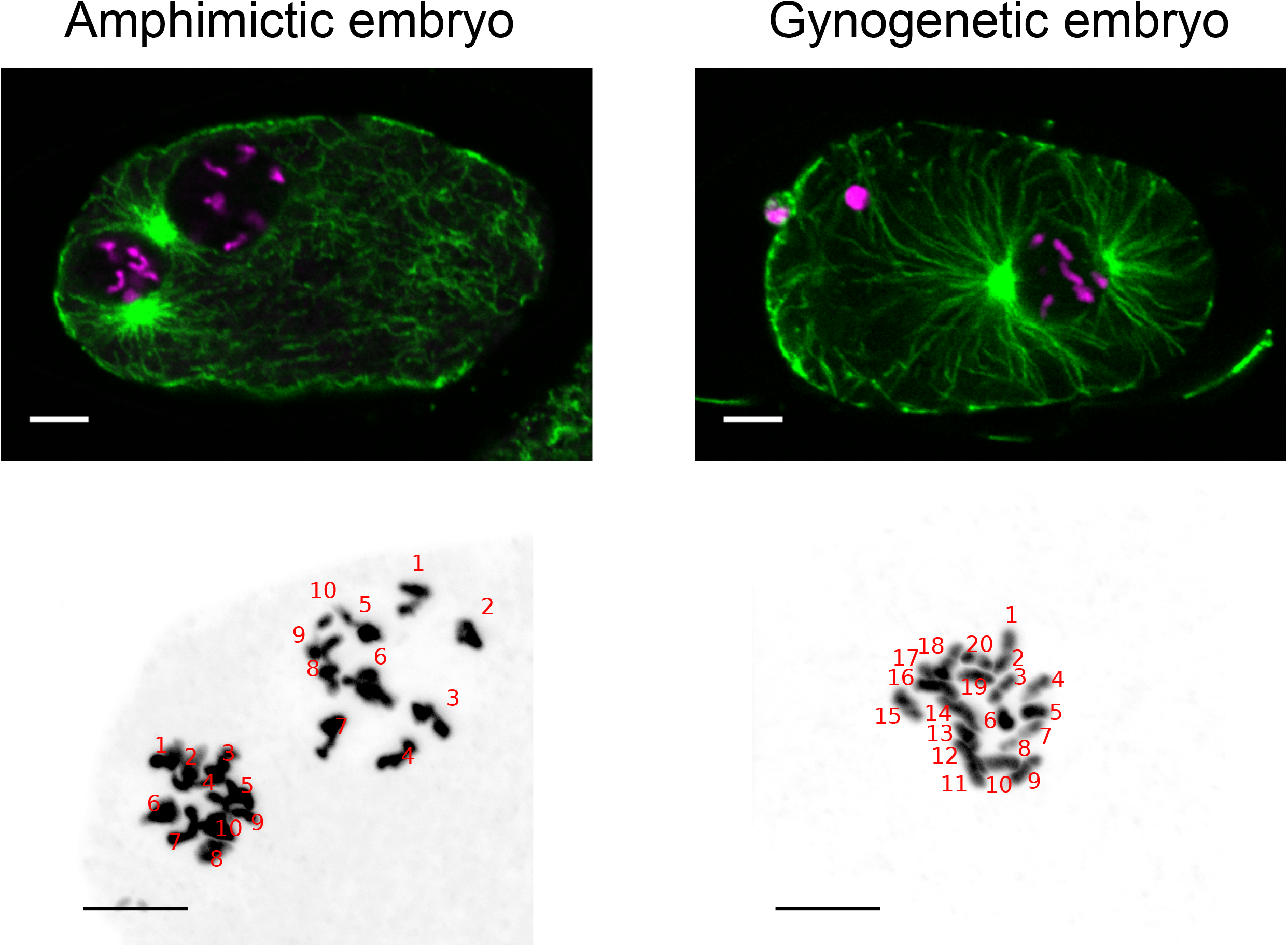
2n=20 chromosomes are found in both types of embryos. The upper panels show a representative amphimictic and a gynogenetic embryo during the reformation of the pronuclei fixed and stained with an anti-tubulin antibody (in green) and with Hoechst to label DNA (in magenta). The lower panel shows a zoom on the chromosomes and a Z projection of the entire nuclei. Chromosomes are shown black. Chromosomes have been numbered arbitrarily. Note that because the amphimixic embryo is slightly younger than the gynogenetic embryo, chromosomes are less condensed and less individualized. Each nucleus of amphimixic embryos show 10 chromosomes, while the single pronucleus of gynogenetic embryos have 20 chromosomes. Scale bar is 5 microns.

**Table S1:**
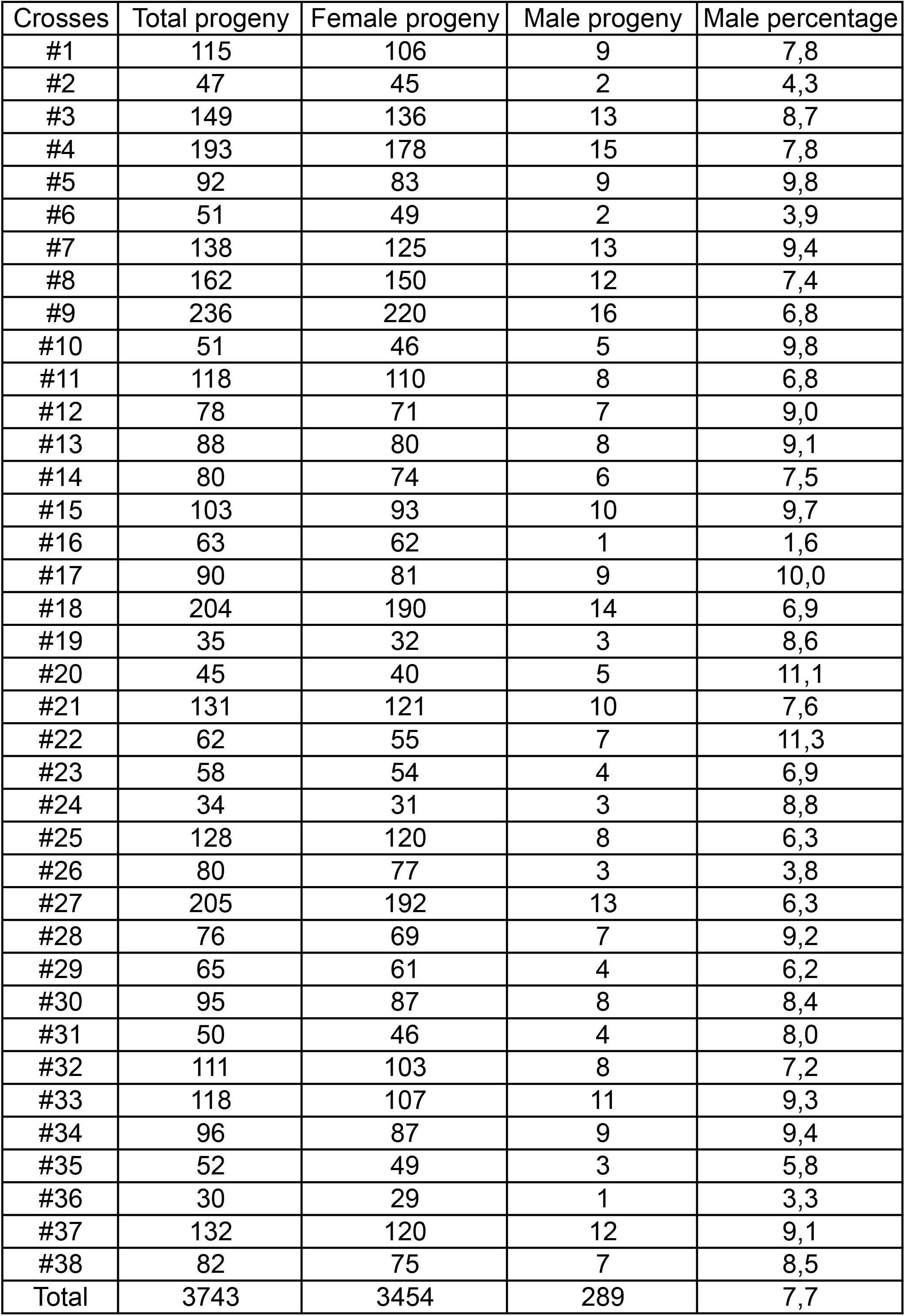
Male percentage in the *M. belari* strain. 38 crosses between 5 virgin females and 2 males were performed on individual plates. The number of males and females F1s was counted when sexes were distinguishable.

**Table S2:** Incidence of the % of males on the total brood size of females. 20 virgin females were isolated on plates and crossed with 1 to 5 males. The total number of embryos laid by the females was counted. Crosses that did not yield any embryo were not taken into account for the calculation of the mean brood size.

| Crosses | 20 ♀ x 1 ♂ |  | 20 ♀ x 2 ♂ |  | 20 ♀ x 3 ♂ |  | 20 ♀ x 4 ♂ |  |
| --- | --- | --- | --- | --- | --- | --- | --- | --- |
| #1 | 1 | 431 | 1 | 366 | 1 | 856 | 1 | 862 |
| #2 | 2 | 819 | 2 | 444 | 2 | 648 | 2 | 552 |
| #3 | 3 | 0 | 3 | 875 | 3 | 697 | 3 | 705 |
| #4 | 4 | 0 | 4 | 769 | 4 | 1132 | 4 | 863 |
| #5 | 5 | 372 | 5 | 1105 | 5 | 461 | 5 | 1023 |
| #6 | 6 | 789 | 6 | 1046 | 6 | 296 | 6 | 603 |
| #7 | 7 | 191 | 7 | 596 | 7 | 800 | 7 | 726 |
| #8 | 8 | 511 | 8 | 671 | 8 | 1261 | 8 | 626 |
| #9 | 9 | 386 | 9 | 950 | 9 | 965 | 9 | 1001 |
| #10 | 10 | 544 | 10 | 272 | 10 | 548 | 10 | 698 |
| #11 | 11 | 158 | 11 | 284 |  |  |  |  |
| #12 | 12 | 0 | 12 | 746 |  |  |  |  |
| #13 | 13 | 709 | 13 | 1346 |  |  |  |  |
| #14 | 14 | 705 | 14 | 1315 |  |  |  |  |
| #15 | 15 | 324 | 15 | 452 |  |  |  |  |
| #16 | 16 | 499 | 16 | 1108 |  |  |  |  |
| #17 | 17 | 534 | 17 | 909 |  |  |  |  |
| #18 | 18 | 0 |  |  |  |  |  |  |
| #19 | 19 | 621 |  |  |  |  |  |  |
| #20 | 20 | 249 |  |  |  |  |  |  |

**Table S3:** Preference for sib-sib mating. Gravid females were isolated on single plates. F1 females were crossed with F1 males from the same mother (sib-sib) or from another female (non siblings crosses). The number of crosses that yielded a progeny (successful cross) was counted. The expriment was repeated 3 times and each time, 50 crosses of each category were performed.

|  | Siblings crosses |  |  | Non siblings crosses |  |  |
| --- | --- | --- | --- | --- | --- | --- |
| Experiment | Total | Successful | %age of succesful crosses | Total | Successful | %age of successful crosses |
| #1 | 50 | 37 | 74 | 50 | 24 | 48 |
| #2 | 50 | 41 | 82 | 50 | 24 | 48 |
| #3 | 50 | 39 | 78 | 50 | 25 | 50 |
| Total | 150 | 117 | 78 | 150 | 73 | 49 |

## Appendix 1: solution of the analytical model

Strategy *x*^*^ is evolutionarily stable when

∀*x* ≠ *x*^*^, *W*(*x*^*^) > *W*(*x*) which implies that *dW*(*x*) / *dx* = 0 when *x* = *x*^*^

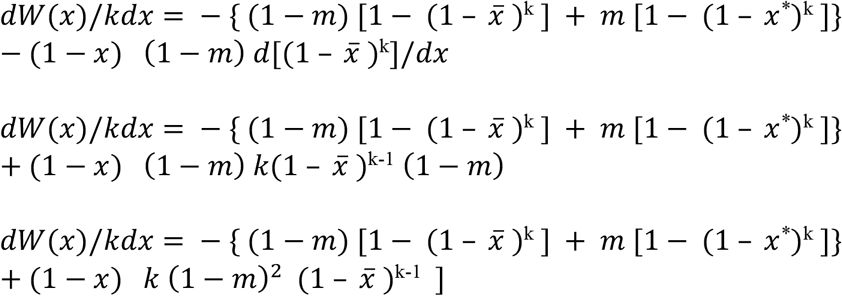

This value must be 0 when *x* = *x*^*^ (note that then, 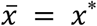)

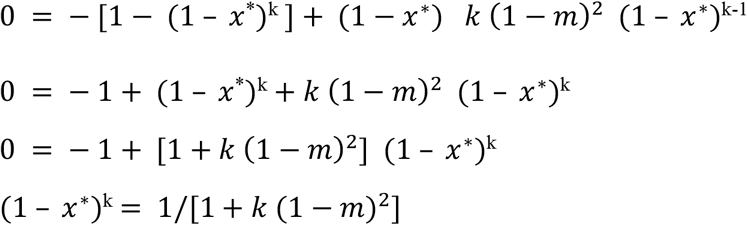

And the ESS proportion of females is thus

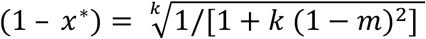

